# Natural and engineered isoforms of the inflammasome adaptor ASC form non-covalent, pH-responsive hydrogels

**DOI:** 10.1101/2023.05.03.539154

**Authors:** Eduardo A. Gaspar-Morales, Anthony Waterston, Pedro Diaz-Parga, Ariell M. Smith, Mourad Sadqi, Arvind Gopinath, Roberto C. Andresen Eguiluz, Eva de Alba

## Abstract

The protein ASC polymerizes into intricate filament networks to assemble the inflammasome, a filamentous multiprotein complex that triggers the inflammatory response. ASC carries two Death Domains integrally involved in protein self-association for filament assembly. We have leveraged this behavior to create non-covalent, pH-responsive hydrogels of full-length, folded ASC by carefully controlling the pH as a critical factor in the polymerization process. We show that natural variants of ASC (ASC isoforms) involved in inflammasome regulation also undergo hydrogelation. To further demonstrate this general capability, we engineered proteins inspired in the ASC structure that successfully form hydrogels. We analyzed the structural network of the natural and engineered protein hydrogels using transmission and scanning electron microscopy, and studied their viscoelastic behavior by shear rheology. Our results reveal one of the very few examples of hydrogels created by the self-assembly of globular proteins and domains in their native conformation and show that Death Domains can be used alone or as building blocks to engineer bioinspired hydrogels.

## INTRODUCTION

Protein- and peptide-based hydrogels are desirable biomaterials for biomedical applications because of their inherent biodegradability, the potential to incorporate a vast variety of functions and their capability to respond to stimuli.^1–6^ Biomedical applications of biological hydrogels include emulating artificial extracellular matrices as scaffolds for tissue engineering, drug delivery by controlled release, biomaterials for wound repair and surgery procedures, biosensors and imaging.^6–8^ In addition, reconstituted protein-based hydrogels are used to study the function and properties of biological hydrogels such as mucus.^9^ Thus, intensive research and bioengineering efforts are currently directed to design and produce new protein-based hydrogels with tunable mechanical properties and specific functionalities.

Most currently known protein-based hydrogels are composed of fibrous proteins, such as collagen, and proteins that are major components of the extracellular matrix, like elastin.^3,10–12^ In contrast, globular proteins in their native state are far less commonly capable to transition to a hydrogel state. However, upon denaturation by acidic pH, high temperature or addition of alcohols, globular proteins unfold acquiring self-assembly capabilities that facilitate hydrogel formation.^5^ In addition, numerous examples have been reported of hydrogels formed by the self-assembly of α-helical peptides into coiled coils,^1,13–15^ and β-sheet forming peptides.^1,5,16^ More recently, a few examples have emerged of hydrogels formed by chemical cross-linkage of globular protein domains using enzyme catalyzed coupling, photo-induced crosslinking, and click-chemistries.^1^ Additionally, globular proteins have been used to create hydrogels with physical non-proteinaceous^1,17,18^ and proteinaceous^1,19–21^ cross-linkages. An elegant example of such hydrogel is based on the high binding affinity between WW domains and polyproline rich motifs.^22^ The design of this all-proteinaceous hydrogel consists in mixing two polypeptides carrying several WW domains and polyproline rich motifs, respectively, spaced by short peptide linkers. The viscoelastic behavior, specifically the storage or shear modulus, of the different protein-based hydrogels depends on the molecular composition and type of cross-linkage. Importantly, it has been demonstrated that the stability of the globular fold strongly influences the hydrogel mechanical response.^17,23–25^ Thus, globular-protein hydrogels may offer a broader scope of applications by increasing elasticity and toughness *via* the manipulation of the unfolding/folding processes of the globular domains that act as the hydrogel building blocks. ^17,23–25^ For instance, protein unfolding will result in the extension of the polymer length that could in turn lead to modifications of the viscoelastic behavior.^25^

Notably, examples of hydrogels solely based on the self-assembly of folded proteins are lacking. Here we show that globular Death Domains^26^ have the capability to form hydrogels without requiring any molecular helper such as peptide-based or synthetic cross-linkers. To demonstrate hydrogelation and study the properties of hydrogels formed by Death Domains, we have used two natural proteins involved in the inflammatory response. Specifically, we have studied hydrogel formation by the natural protein ASC^27–29^ and its isoform ASCc,^30^ which activate and inhibit the inflammatory response, respectively.^30^ The three-dimensional structure of ASC reveals that the protein is composed of two Death Domains, PYD (pyrin) and CARD (caspase activation and recruitment domain), connected by a linker.^28,29,31^ No structural information is known about ASCc; however, its amino acid sequence suggests an intact CARD connected to an incomplete PYD. Finally, to test whether this behavior is limited to natural Death Domain proteins, we have engineered constructs based on the ASC structure to analyze the hydrogelation capabilities of non-natural proteins.

## MATERIALS AND METHODS

### Protein Expression and Purification

Plasmids (pET-15b) encoding the sequences of ASC, ASCc, CARD and CARD-CARD (Supporting Figure S1) were transformed in BL21(DE3) *E. coli* cells. Bacteria were grown in LB medium overnight at 37 °C. Cell cultures were diluted to reach an OD of 0.8 at 600 nm and were subsequently induced for 4 hours by the addition of 1 mM IPTG to produce overexpression of the recombinant proteins. Cells were harvested by centrifugation at 8,000 rpm for 30 minutes at 4 °C. The pellets were resuspended in 20 mM Tris-HCl pH 8, 500 mM NaCl, 5 mM imidazole, and 6 M guanidinium hydrochloride (resuspension buffer) and then sonicated (twelve times in 15-second intervals, alternated with 45-second resting periods). The lysed cells were ultracentrifuged at 35,000 rpm for 30 minutes at 4 °C. The resulting supernatant was filtered to remove cell debris with a 0.45 µm pore-size filter. All proteins have a 6-histidine tag at the N-terminus and were purified by nickel affinity chromatography. The chromatography matrix was equilibrated with the resuspension buffer. After protein binding, the matrix was washed with a buffer containing 20 mM Tris-HCl pH 8, 500 mM NaCl, 20 mM imidazole and 6 M guanidinium hydrochloride. The purified proteins were eluted in a 45-minute gradient with a buffer containing 20 mM Tris-HCl pH 8, 500 mM NaCl, 500 mM imidazole, and 6 M guanidinium hydrochloride. The solutions of the eluted proteins were dialyzed against 1 L of a dialysis buffer containing 0.5 M TCEP at pH 3.8 using dialysis cassettes, and the buffer was changed 3 times every 2-3 hours. The protein solutions were extracted from the dialysis cassette and filtered. All proteins were further purified by reverse phase chromatography in a C4 column. The C4 matrix was equilibrated with a buffer containing 94.9% H_2_O, 5% acetonitrile, 0.1% trifluoroacetic acid, and the proteins were eluted in a 30 – 40-minute gradient with an elution buffer containing 94.9% acetonitrile, 5% H_2_O, 0.1% trifluoroacetic acid. All eluted protein solutions were lyophilized for the removal of organic solvents and stored in a desiccator. All buffers were filtered through 0.2 μm pore-size filters.

### Peptide synthesis for the formation of the CARD-peptide hydrogel

The sequence of the peptide from N- to C-termini is MGRARDAILDALENLTAEELKKFKLQAAT, which encompasses the incomplete PYD domain of ASCc (Supporting Figure S1). This peptide, with no modifications at the N- or C-termini, was synthesized by solid-phase peptide synthesis by Thermo-Fisher with a purity > 95%.

### Mass spectrometry

The purity and integrity of the recombinant proteins were determined by mass spectrometry and SDS-PAGE. Lyophilized protein material was dissolved in a solution containing 95% acetonitrile, 4.9% water, 0.1% formic acid and injected into an electrospray ionization mass spectrometer (Q-Exactive Hybrid Quadrupole-Orbitrap, Thermo). The molecular weight obtained by mass spectrometry matched the expected molecular weight based on the amino acid sequence for all proteins.

### Hydrogel Formation

The purified proteins were dissolved in aqueous solution containing 500 μM TCEP at pH 3.8. The protein concentration was determined by absorbance spectroscopy using the Lambert-Beer law at 280 nm using the theoretical extinction coefficients predicted by the Expasy server^32^ based on amino acid sequences. We use lyophilized protein at ≥ 90 - 95% purity determined by mass spectrometry and polyacrylamide gel electrophoresis.

Hydrogelation started from solutions of ASCc and CARD-peptide at 0.3 mM, and ASC and CARD-CARD at 0.5 mM and 0.9 mM, respectively. All hydrogels were formed by slowly increasing the pH from 3.8 to 6.8. Basification was done by stepwise addition of small volumes (2.5 – 4 µL) of dilute solutions of NaOH (0.1 M) to the concentrated protein solutions. Incubation periods of 5 to 10 minutes followed each basification step to allow the formation of the complex filament networks over time. The slow basification process starts at pH 3.8 once the CARD and PYD domains are properly folded.^27^ The small volumes of the NaOH solution must be pipetted into the center of the protein solution for an even distribution. The solutions are swirled and set aside for 5 minutes. The pH is monitored after each incubation time with a microelectrode until the pH value reaches 4.8. Subsequent pH increase is achieved by adding approximately 2 µL of 0.05 M NaOH. The flowability of the protein solution decreases at pH > 5. At this point, the incubation time increases from 5 to 10 minutes. The solutions are gently swirled to prevent disruption of the hydrogel network up to the required pH of 6.8. The hydrogelation process for ASC is slower to prevent precipitation. For ASC hydrogelation, the pH is increased at a rate of 0.5 pH units every 120 minutes. Changes in the proteins electrostatic surfaces are not expected in the pH range from 6.8 to 8.3 as no amino acids titrate in this pH interval. In addition, pH values higher that 8.3 (titration of the sulfhydryl in the cysteine side chain) could lead to protein unfolding. Thus, hydrogel formation and behavior were not studied at pH > 6.8.

### Swelling Ratio

Hydrogel swelling ratios were obtained by lyophilization using equation 1.

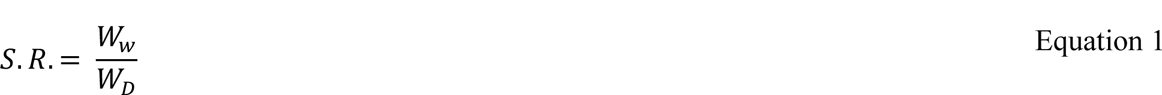

Where *S. R.* is the swelling ratio, *W_w_* and *W_D_* is the weight of the wet and dry hydrogel, respectively.

The hydrogel formed in a glass vial is weighed, then flash-frozen in liquid nitrogen followed by lyophilization. The weight of the wet (*W_w_*) and dry (*W_D_*) hydrogel is obtained before and after lyophilization, respectively, by subtracting the weight of the empty lyophilized vial.

### Transmission Electron Microscopy

A volume of 4 µl of the hydrogel was deposited on a 300 mesh Cu TEM grid that had been glow discharged previously. After 10-minute incubation, the grid was washed for 10 seconds in three 40 µl droplets of HPLC water and stained in three 40 µl droplets of 2% uranyl acetate. After staining the grid for 5 minutes, the excess of staining solution was wipe dried. Images of the hydrogels were obtained using a Talos F200C G2 Transmission Electron Microscope with a Field emission Gun operating at 200 kV. Images were captured with a Ceta 16M Camera. The open-source software ImageJ with the options “analyze” and “measure” was used to analyze the dimensions of hydrogel filaments and fibrils.

### Scanning Electron Microscopy

On an aluminum specimen mount, lyophilized protein was deposited on conductive carbon adhesive tape. Samples were coated for 40 seconds with a gold sputter coater at a vacuum of 0.07 torr, corresponding to approximately 5 nm gold film. Images were obtained using a Zeiss Gemini 500 Field Emission Scanning Electron Microscope operating at voltage of 3 kV. The open-source software ImageJ was used for analysis. The images were first inverted and then the option “analyze particles” and “measure” were used to determine hydrogel pore dimensions with a threshold of 2 μm^2^ and no upper limit.

### Rheology Experiments

Bulk shear rheological behaviors of ASC, ASCc, CARD-peptide, and CARD-CARD hydrogels at pH 6.8 were investigated using an MCR-302 (Anton Paar) at 25 °C. We used a stainless-steel parallel plate attachment (Anton Paar PP-25, 25 mm diameter) for all experiments. The hydrogels were kept at ∼ 4 °C and allowed to reach room temperature (∼ 25 °C) for 45 minutes. A total of 300 μL of previously crosslinked samples were then carefully taken from a glass vial and loaded onto the center of the bottom steel plate using a plastic spatula. Once in place, the stainless-steel top parallel plate was carefully and slowly lowered until reaching a working thickness of 500 μm and filling the parallel plate cavity. In this process, it was ensured that the sample network was still intact. The storage modulus (G’) and loss modulus (G’’) were then measured as a function of shear strain, γ, and frequency, ω. The storage or the shear modulus G’ relates the shear stress that is in phase with the shear strain and provides a measure of the elasticity of the hydrogel network as a function of the forcing frequency or the forcing shear strain. The loss modulus G’’ is a measure of the effective dissipation due to viscosity in the hydrogel and manifests as a phase lag between shear strain and the shear stress. Tests were conducted with a humidity chamber to minimize solvent evaporation from the hydrogel samples.

The storage and loss moduli were independently measured using frequency sweeps and shear strain sweeps. The frequency dependence of G’ and G’’ was investigated by imposing the sample to small amplitude oscillatory frequency. The angular frequency, ω, varied between 0.1 to 200 rad/s. The amplitude of the shear strain γ was held constant at 1%, consistent with previously published parameters.^33^ Next, the strain response of G’ and G’’ was then investigated by subjecting new samples to oscillatory shear at constant angular frequency ω of 6.28 rad/s (equivalent to 1 Hz) while varying the strain amplitude γ, from 0.1 to 100 %.^33^ Two independent experiments for frequency and strain responses were performed per hydrogel, corresponding to 16 hydrogel samples.

## RESULTS

### Protein design for hydrogel formation based on the structure and oligomerization properties of ASC

Previously, we showed that full-length ASC polymerizes into filaments and filament bundles *via* homotypic protein-protein interactions mediated by its two oligomerization domains, PYD and CARD.^31,34^ Based on the analysis of TEM micrographs, we reported that these filaments are ∼ 7 nm wide and reach lengths of 1 μm.^31,34,35^ The filaments assemble laterally forming bundles of 2- to 7-filaments. Oligomerization is not restricted to the full-length protein, as we have shown that ASC’s individual domains are capable of polymerizing into filaments as well.^34,36,37^ Using solution NMR, we found that homotypic CARD-CARD binding affinity is slightly higher than that of PYD-PYD binding.^34,36^ We also demonstrated that ASC interactions are pH dependent. ASC is properly folded and monomeric at pH = 3.8; however, it precipitates at higher pH values.^27^ The 3D structure of ASC (Figure 1A) reveals the protein’s electrostatic surface composed of positively and negatively charged patches, as well as some non-polar regions involved in ASC filament formation.^27^ At low pH, the negative charges are neutralized, thus shielding the electrostatic interactions and shifting the monomer-oligomer equilibrium towards the monomeric species. Upon pH increase, deprotonation leads to a larger number of negative charges leading to favorable intermolecular electrostatic interactions that are essential for the non-covalent polymerization of the protein.^34^

**Figure 1.**
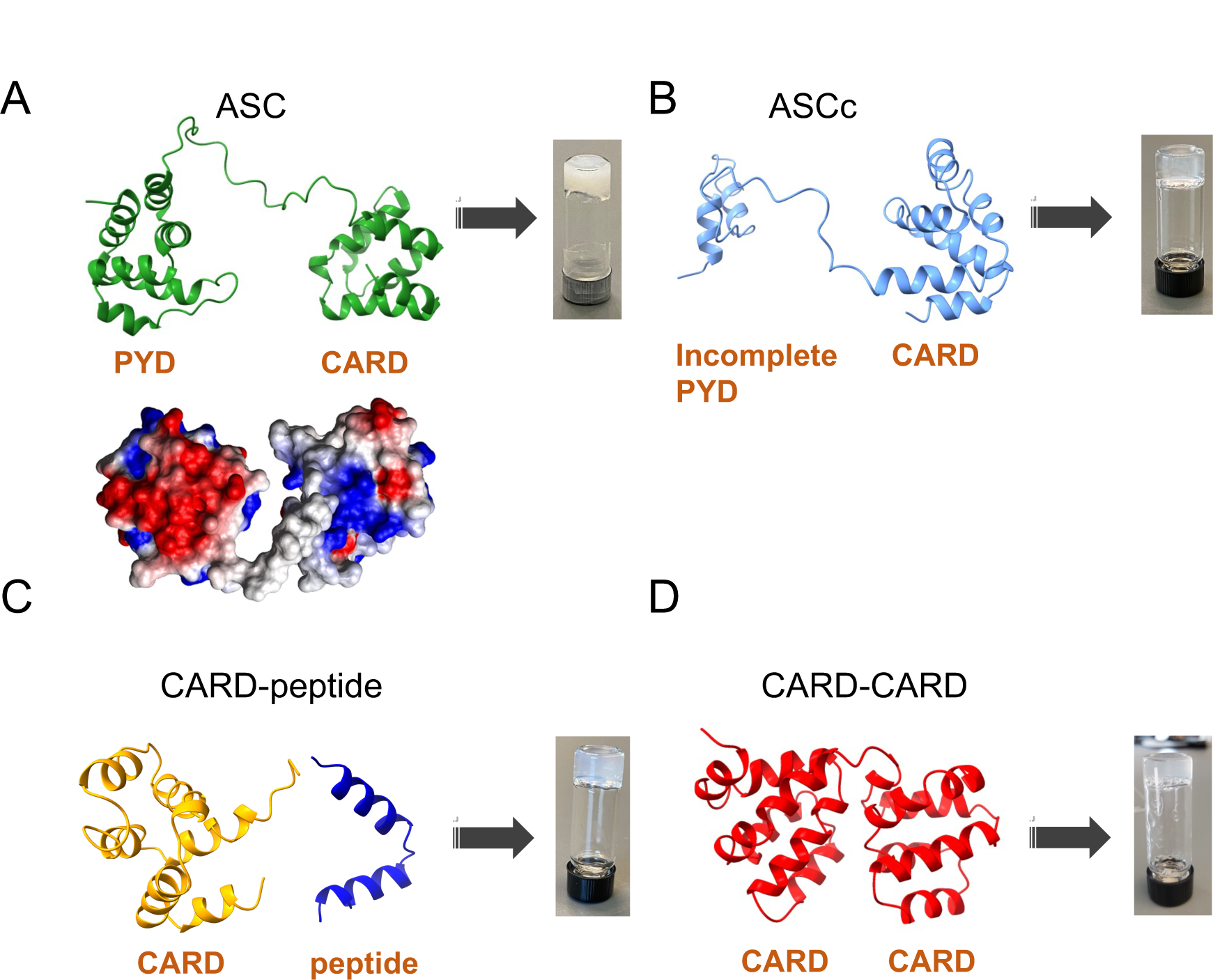
Three-dimensional structures of the proteins used for hydrogel formation. A) ASC experimental structure determined by solution NMR.^29^ Top: ribbon diagram depicting the polypeptide backbone; bottom: electrostatic surface showing negatively and positively charged patches in red and blue, respectively; B) ASCc structural model; C) CARD-peptide model: CARD structure from ASC^29^ and peptide structural model; D) CARD-CARD model structure. Images of the respective hydrogels formed by these proteins are shown next to each model structure. The structural models were created with I-Tasser.^40^ The ribbon diagrams of the proteins’ backbone were done with ChimeraX.^41^

Recently, a short peptide inspired by the amino acid sequence of the ASC-PYD domain with the addition of the self-assembly moiety Nap-FF (a naphthalene aromatic ring followed by two phenylalanine amino acids) has been shown to form nanofibers.^38^ The addition of vitamin B6, which has been suggested to influence the self-assembly of ASC,^38,39^ is needed for the hydrogelation of this short peptide. The peptide’s sequence is Nap-FF-KKFKLKL.^38^ In an analogous work, this positively charged peptide has been shown to form hydrogels upon mixing with a peptide of complementary charge also inspired in ASC-PYD.^33^ The peptide sequences of the negative counterpart are Ac-DALDLTD or Nap-FF-DALDLTD.^33^ The presence of at least one self-assembly moiety (Nap-FF) in the positively or the negatively charged peptides is required for hydrogelation.

Based on these studies and considering the polymerization capabilities of ASC and its individual domains, we hypothesized that full-length ASC could also form hydrogels. This is an important question because there are almost no examples of globular proteins/domains in their native conformation that are known to form hydrogels. To test our hypothesis, we aimed at creating hydrogels solely composed of natural ASC. In addition, ASC has several isoforms with similar structural characteristics.^30^ Thus, we speculated whether the isoforms could also form hydrogels. We specifically focused on the isoform ASCc that has been shown to inhibit, rather than activate the inflammasome.^30^ The inhibition mechanism is not known, but it has been postulated that ASCc might interfere with ASC self-assembly thus disrupting the formation of the inflammasome.^30^ To gain understanding on the conformation of ASCc, we created a structural model with the program I-Tasser that uses a database of experimental 3D protein structures.^40^ The model indicates that ASCc likely carries an intact CARD and an incomplete PYD (Figure 1B). ASCc will likely polymerize despite the incomplete PYD since we demonstrated that ASC-CARD polymerizes into filaments^34^ and the amino acid sequence of ASCc-CARD shares 100% identity with ASC-CARD.^30^ To determine the role of the incomplete PYD in the potential hydrogelation of ASCc, we created an artificial protein carrying ASCc-CARD and attempted hydrogelation in the presence and absence of a synthetic peptide comprising the residual PYD (Figure 1C). Finally, we engineered a completely artificial protein carrying two CARD domains in tandem to study hydrogelation (Figure 1D). We chose the CARD domain due to the stronger affinity for homo-oligomerization.^34,36^ The names we will use henceforth for these ASC-inspired proteins refer to their respective molecular compositions: ASC, ASCc, CARD-peptide, CARD-CARD (Figure 1).

### The hydrogelation process of ASC-inspired proteins is pH-dependent

ASC, ASCc as well as the PYD and CARD domains oligomerize and precipitate when the solution pH is increased quickly from acidic to neutral values. Our data show that massive precipitation occurs by pH increase in the 0.3 – 1 mM protein concentration range. However, a slow and tightly controlled basification leads to the formation of hydrogels (Figure 2). The 3D NMR structure of ASC was determined at pH 3.8 demonstrating that the protein is properly folded.^29,37^ Thus, the hydrogelation process must start at this pH to ensure the formation of the correct protein electrostatic surface required for proper oligomerization. The pH value is gradually increased to pH 4.8 in three to four 5-minute steps. Basification is set at a significantly slower pace after pH 4.8, requiring multiple steps spaced by 10 minutes. The slow pH-increase process for hydrogelation takes approximately 4 hours.

**Figure 2.**
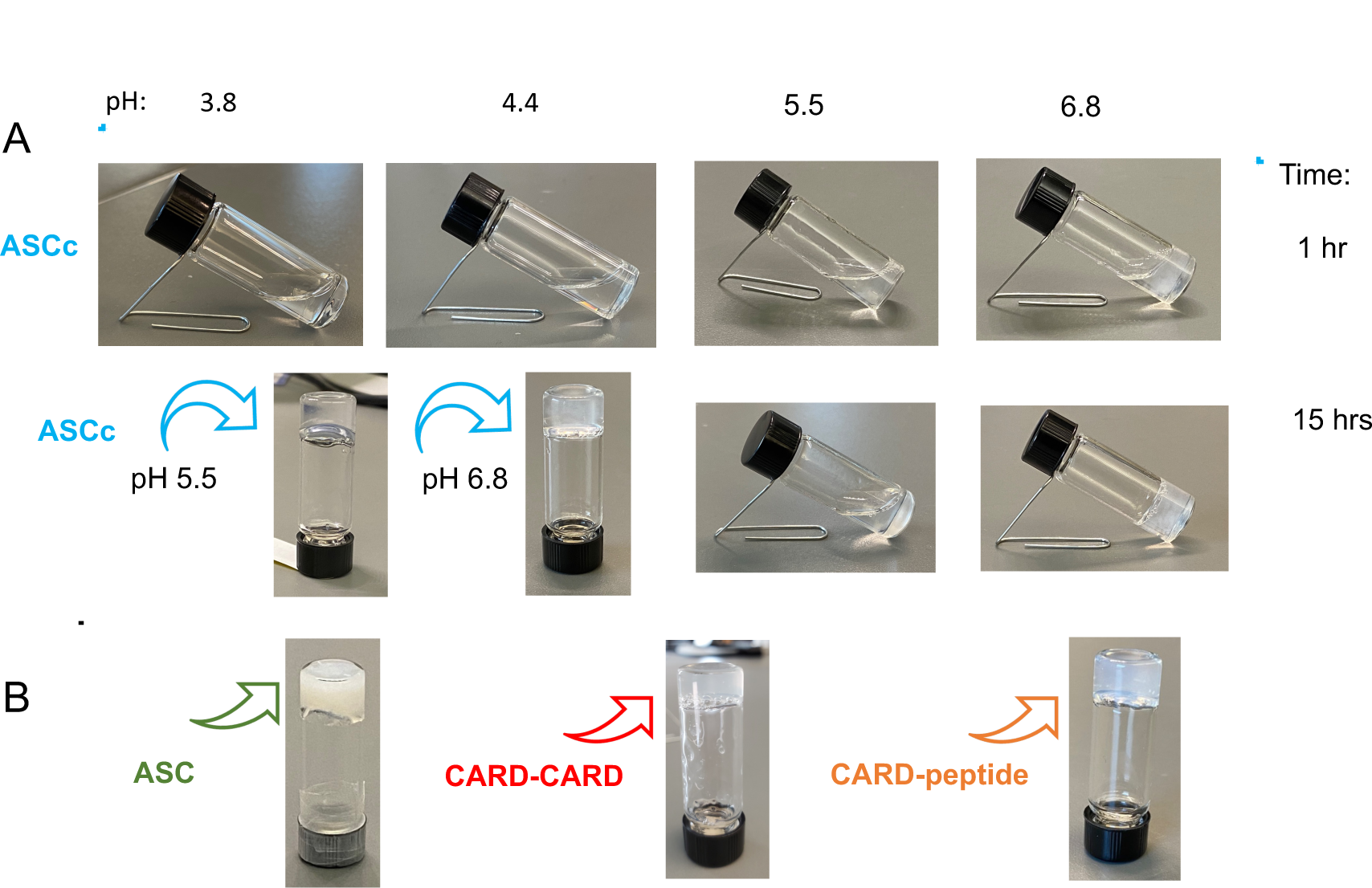
pH-dependent hydrogelation of ASC, ASCc, CARD-peptide and CARD-CARD. A) Hydrogelation process of ASCc as a function of pH increase and time. B) Hydrogels formed by ASC, CARD-CARD and CARD-peptide by analogous pH increase and overnight incubation.

Upon pH increase to 6.8, the protein solutions thicken, suggesting increased viscoelastic characteristics and finally resulting in hydrogel formation upon overnight incubation at room temperature. The first indication of hydrogelation is the decreased flowability and solid-like appearance of the solution, which fails to flow to the bottom of the vial when inverted (Figure 2). Reaching a pH of 6.8 is important for hydrogel consistency, as overnight incubation at more acidic pH (e.g., pH 5.5) leads to more flowable hydrogels (Figure 2A). This result points to the importance of the negatively charged regions of the protein surface for polymerization and hydrogel formation, resulting from the deprotonation at pH values higher than 5.5. Overnight hydrogelation was obtained for ASC and ASCc (0.5 mM and 0.3 mM protein concentration, respectively), which implies that the complete PYD (Figure 1) is not required and suggests that hydrogelation might be possible for constructs containing only the CARD domain.

Thus, we engineered a CARD-only construct (no PYD or incomplete PYD linked) and tested its hydrogelation capabilities. The CARD-only protein solution (0.3 mM) thickened upon slow pH-increase after overnight incubation (Supporting Figure S2). Additional incubation time and basification was required for hydrogelation. In contrast, increasing the concentration of the CARD-only protein solution to 0.5 mM did not improve the hydrogelation capabilities. The CARD-only hydrogel showed more flowability compared to ASC and ASCc hydrogels as it deforms when inverted (Supporting Figure S2). This was a surprising result as ASCc with a CARD and an incomplete PYD is capable of overnight hydrogelation. Therefore, we tried to emulate this structure by mixing equimolar concentrations of the CARD-only protein with a synthetic peptide comprising the incomplete PYD domain of ASCc (hydrogel: CARD-peptide at 0.3 mM) (Figure 1C). This heterotypic interaction facilitated overnight hydrogel formation of the CARD-only construct with the synthetic peptide (Figure 2B). We also tested the behavior of CARD-only in the presence of minor amounts of the peptide, instead of equimolar concentration, observing that hydrogelation is not complete after overnight incubation, but the hydrogel eventually forms after several days. This result indicates that equimolar amounts of the peptide emulating the incomplete PYD promote hydrogelation. The CARD-peptide hydrogel is an example of the possibility to form a hydrogel with two separated molecular components that are otherwise linked in the natural ASCc isoform. However, the hydrogelation process is significantly perturbed when one of the components is missing. The capability of CARD-only and CARD-peptide to form hydrogels indicates that the linker connecting the two domains in ASC and ASCc (Figure 1) is not essential for hydrogel formation. However, it has been demonstrated that the linker tethering the two Death Domains plays an important role in the initial stages of the oligomerization kinetics of ASC isoforms.^31,35^

The CARD-peptide hydrogel is an artificial construct designed based on the structure of ASCc (i.e., the CARD of ASCc and a synthetic peptide comprising the amino acids sequence of the incomplete PYD) (Figure 1). To test whether another artificial construct could form a hydrogel, we engineered the protein CARD-CARD, with two CARD domains in tandem connected by a short linker. The CARD domain was selected for this design to maximize hydrogelation capabilities as the CARD shows slightly higher affinity for self-association than the PYD.^34^ The CARD-CARD protein (0.9 mM) is also capable of hydrogelation following an analogous process of pH increase and overnight incubation (Figure 2B). This result suggests that the PYD can be replaced by an additional CARD for protein hydrogelation.

It is important to note that the isoelectric point (pI) of the proteins is close to neutral pH, and thus very similar to the final pH value for optimum hydrogelation. The pIs are 6.80 (ASC), 7.14 (ASCc), 7.14/6.04 for CARD and peptide, respectively, and 6.83 (CARD-CARD). As commonly observed in globular proteins, the number of positively and negatively charged amino acids (Arg, Lys and Asp, Glu, respectively) at neutral pH in the proteins studied is very similar. The positive and negative charges, respectively, are 23, 24 (ASC); 16, 16 (ASCc); 11, 11 and 5, 5 (CARD-peptide) and 22, 21 (CARD-CARD). Thus, the pH dependence of the hydrogelation process is not related to the number of charges *per se*, but to the clustering of these charges in the protein electrostatic surface (Figure 1A). Nonetheless, we reported previously that hydrophobic interactions also play an important role in ASC self-association.^36^

All four hydrogels are relatively soft and can be easily deformed with a pipette tip or a spatula. This is an expected result as ASC-inspired hydrogels are assembled by physical cross-linkages resulting from electrostatic interactions and from the hydrophobic effect. These weak interactions explain why the hydrogelation process is reversible. Decreasing the pH back to 3.8 regenerates the liquid protein solution. Upon acidification, the negatively charged amino acids are neutralized, thus shielding the electrostatic interactions with the positively charged regions of the protein surface, leading to oligomer dissociation.

ASC-inspired hydrogels are not completely transparent, showing different degrees of opaqueness with the ASC hydrogel being opaque white. The various shades of white suggest different levels of reflected and scattered light, which could be related to the structural characteristics of the hydrogels. Slight precipitation of protein material embedded in the hydrogels could cause haziness as well. To better understand the structure of ASC-inspired hydrogels we used transmission (TEM) and scanning (SEM) electron microscopy.

### The structural scaffold of ASC-inspired hydrogels is formed by an intricate network of protein filaments and fibrils

We have previously shown that ASC and other isoforms polymerize into filaments and filament-bundles in the absence of hydrogelation.^31,34,35^ Therefore, this type of macrostructure is expected to be the essential structural unit of ASC-inspired hydrogels. However, the hydrogelation process could result in modification of these filaments, thus requiring additional analysis by TEM. Because sample thickness is key to obtaining high-quality images, we inserted a micropipette in formed hydrogels to extract a fraction of the hydrogel material. The TEM images of the 4 hydrogels reveal structural units formed by protein filaments and filament-bundles (fibrils) and show that all hydrogels form large, entangled filament networks (Figure 3).

**Figure 3.**
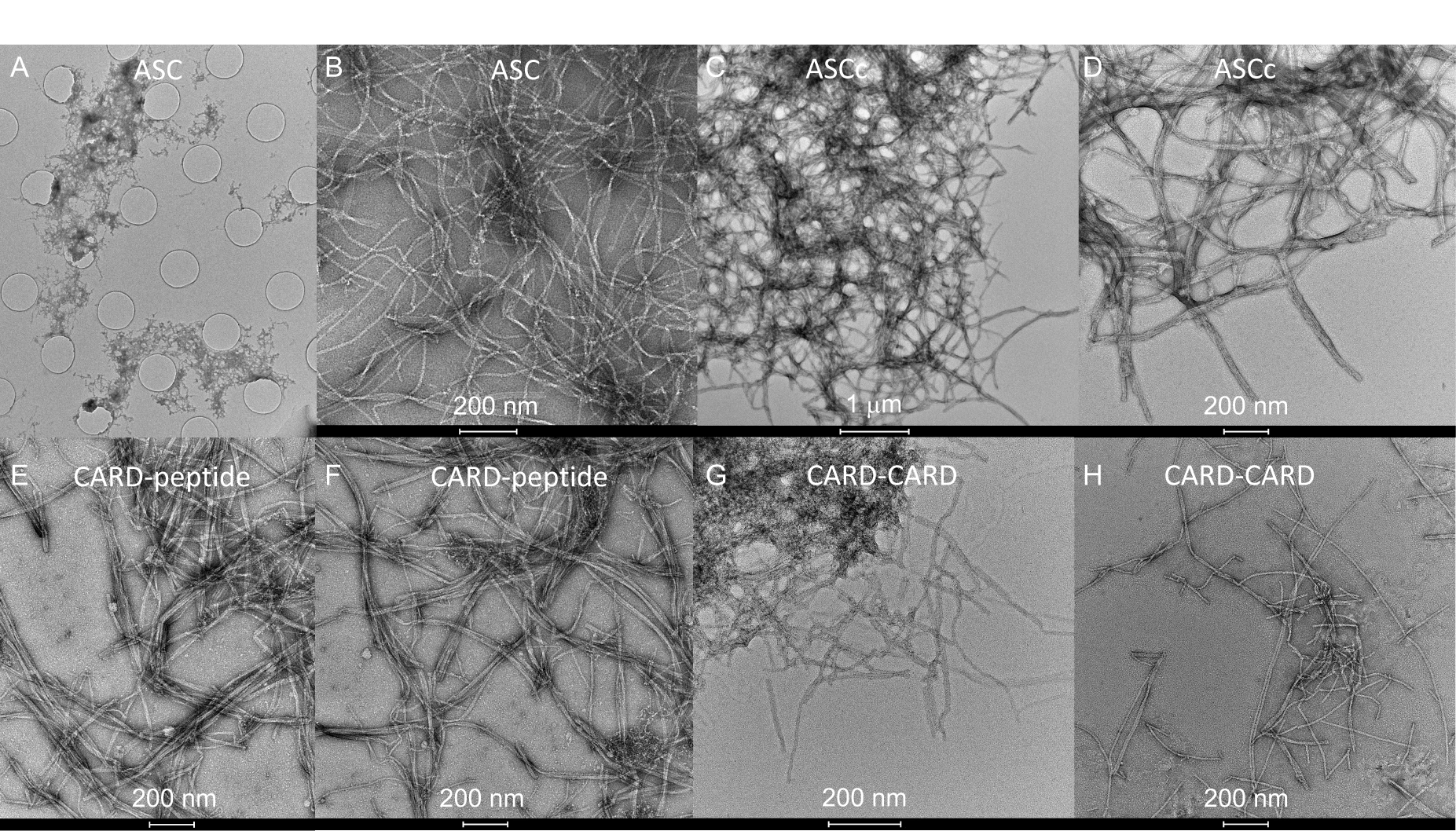
An intricate network of protein filaments and fibrils form the structural scaffold of ASC-inspired hydrogels. Transmission electron micrographs of ASC (A, B), ASCc (C, D), CARD-peptide (E, F) and CARD-CARD (G, H) hydrogels. Scale bar dimensions are indicated. A TEM grid hole size of 1.2 μm in diameter serves as dimension reference for panel A.

These filaments and fibrils were further analyzed to identify potential differences in their dimensions. The analysis of 50 individual filaments per hydrogel indicates that the filaments formed by the different hydrogels have similar widths falling in the 6.1 – 7.9 nm range (Table 1). However, small differences were observed in the frequency of the number of filaments forming fibrils. The analysis of a total of 122 fibrils shows that ASCc and CARD-CARD hydrogels tend to self-assemble into thick bundles composed of 6 and 7 filaments (Figure 4).

**Figure 4.**
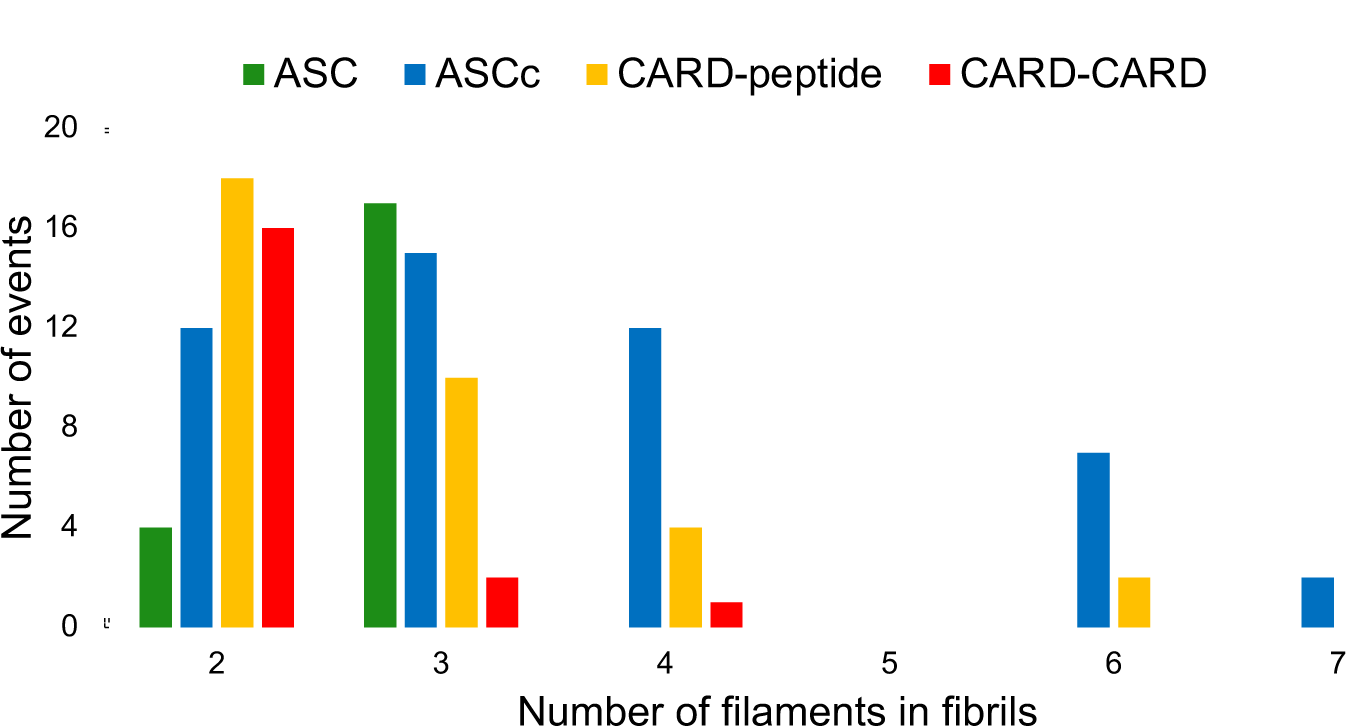
Number of filaments in fibrils formed by ASC-inspired hydrogels. ASCc and CARD-peptide show slightly higher tendency to form filament-bundles composed of 6-7 filaments. Bar height indicates the number of fibrils with the corresponding number of stacked filaments (X axis). Color code is indicated at the top of the figure.

**Table 1.** Analysis of Filaments and Fibrils formed by ASC-inspired hydrogels*.

|  | Hydrogel |  |  |  |
| --- | --- | --- | --- | --- |
|  | ASC | ASCc | CARD-peptide | CARD-CARD |
| Filament width | $7.2 \pm 0.5$ nm | $7.0 \pm 0.9$ nm | $7.3 \pm 0.5$ nm | $7.4 \pm 0.5$ nm |

\* Data from a total of 50 filaments per hydrogel.

### ASC-inspired hydrogels have porous structures

To characterize the macrostructures of the ASC-inspired hydrogels, we have used SEM. Micrographs obtained at different magnifications show that the 4 hydrogels have porous structures, suggesting significantly permeable hydrogels (Figure 5). The surfaces of ASC, CARD-peptide and CARD-CARD appear flaky, whereas ASCc’s surface is more rounded.

**Figure 5.**
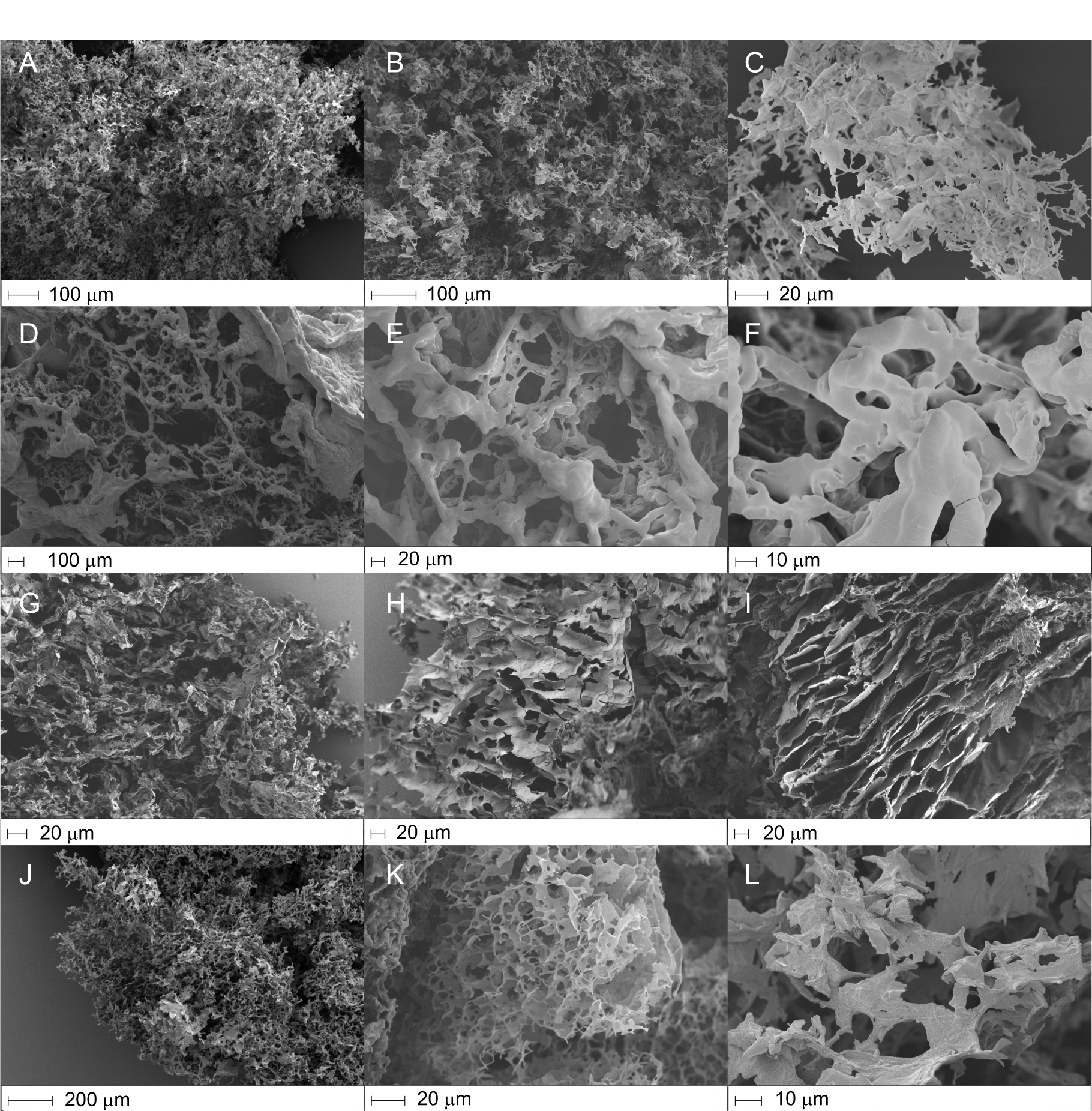
ASC-inspired hydrogels have porous structures. Scanning electron micrographs at different magnification of ASC (A, B, C), ASCc (D, E, F), CARD-peptide (G, H, I), and CARD-CARD (J, K, L) hydrogels. Scale bar dimensions are indicated.

**Figure 6.**
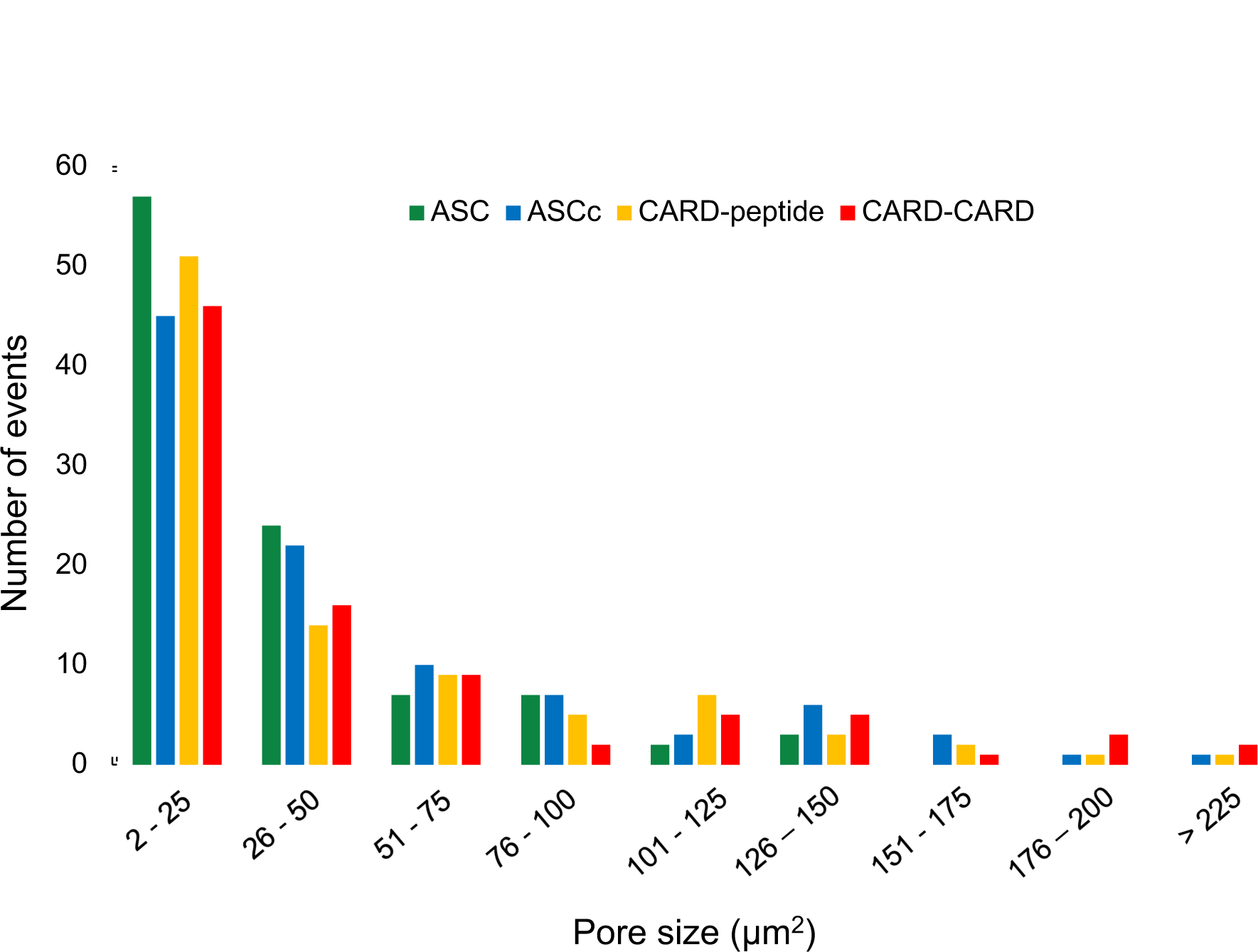
Pore size analysis of ASC-inspired hydrogels. Bar height represents the number of pores with the corresponding area (X axis). Color code is indicated at the top of the figure.

Using these images, we have performed a pore size analysis to identify similitudes and differences between the different hydrogels. The pore size reported here reflects an effective void size and is therefore related to the effective permeability of the hydrogel. The analysis of 380 pores up to 225 μm^2^ shows that the 4 hydrogels tend to form pores smaller than 25 μm^2^, pores sizes in the 25 μm^2^ - 50 μm^2^ rage are also abundant, although less prominent, and sizes larger than 100 μm^2^ are increasingly infrequent. ASC is the only hydrogel for which pores in the 150 μm^2^ - 225 μm^2^ range were not observed.

To connect the pore size distributions with the capability to swell in the presence of water, we measured the hydrogel swelling ratios using lyophilization. The swelling ratios of ASC-inspired hydrogels fall in the range 51 to 123 (Table 2). These values are in accord with physically cross-linked hydrogels that typically do not increase their volume as much as chemically formed hydrogels.^5^ ASC and CARD-CARD show smaller swelling ratios likely due to the higher protein concentration used to form these hydrogels. Correcting these values by considering the difference in concentration compared to ASCc and CARD-peptide results in swelling ratios for ASC and CARD-CARD of ∼ 132 and ∼ 154, respectively. In addition, the precision in the swelling ratio measurements varies significantly from 0.2 to 33 with smaller values for ASC and CARD-CARD. Overall, ASC-inspired hydrogels do not show significant differences in their capacity to absorb water.

**Table 2.**
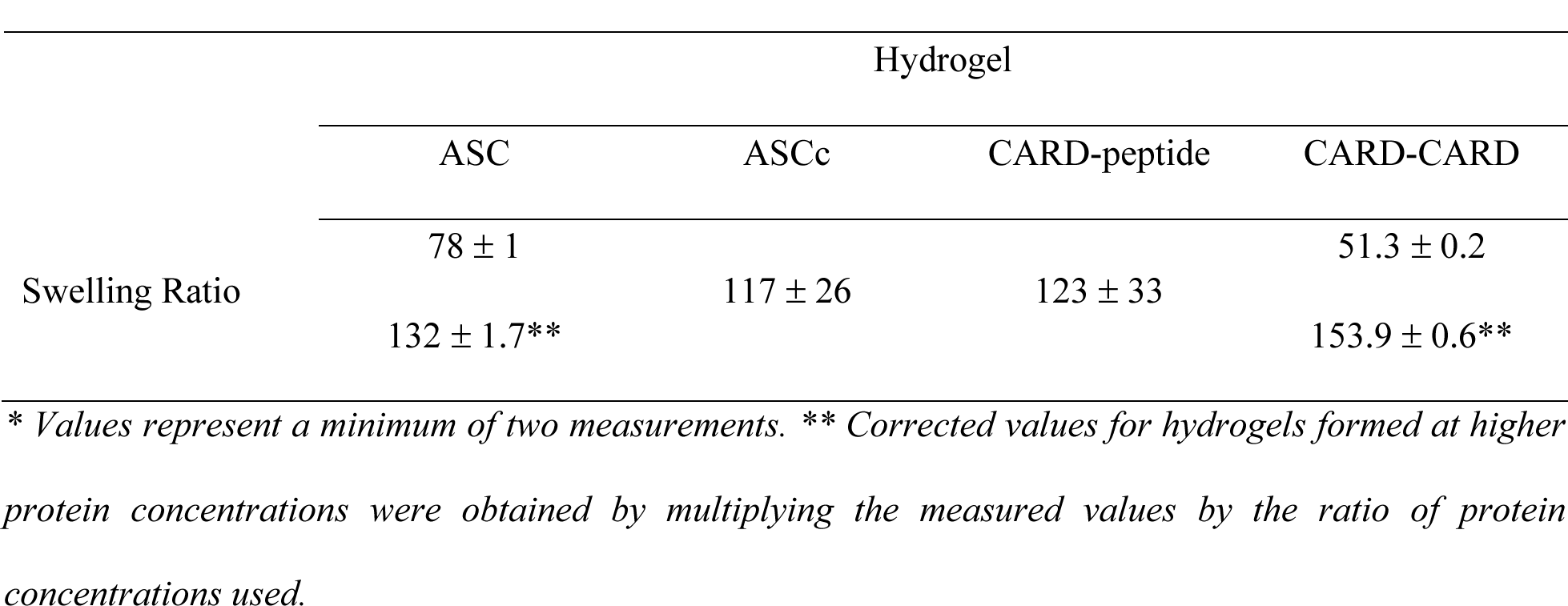
Swelling ratios of ASC-inspired hydrogels*.

### Viscoelastic properties of ASC-inspired hydrogels

To characterize the viscoelastic response of ASC, ASCc, CARD-peptide and CARD-CARD at pH 6.8, we conducted shear rheology measurements. The log-log curves of the storage modulus (G’) and loss modulus (G’’) as a function of angular frequency (*ω*) and shear strain (*γ*) are shown in Figure 7 and Figure 8, respectively. The larger values of G’ compared to G’’ indicate the presence of stable hydrogels with some ability to withstand shear deformations. For the experiments conducted at various values of the angular frequency, our results reveal that the G’ and G’’ values slightly increase with increasing frequency within the frequency range tested, indicating that the hydrogels become stiffer. The shear strain sweep tests show different rheological responses of the hydrogels. All hydrogels show a linear viscoelastic region up to shear strains of approximately 10%, indicating a predominantly elastic response without stiffening. The characteristic strain at the crossover point where the two moduli are approximately the same, quantifies the value at which the rheological response of the hydrogel transitions from predominantly elastic behavior to predominantly liquid-like viscous behavior. We find that this characteristic strain is construct dependent. Specifically, the crossover strains for ASC, ASCc, CARD-peptide, and CARD-CARD are 7.5 Pa @ 15.8 %, 3.4 Pa @ 39.7%, 17.8 Pa @ 19.9 %, and 47.1 Pa @ 15.8 %, respectively. These different crossover points indicate that ASCc behaves as a solid-like hydrogel predominantly over larger shear strain values, approximately twice the strain when compared to ASC, CARD-peptide, and CARD-CARD.

**Figure 7.**
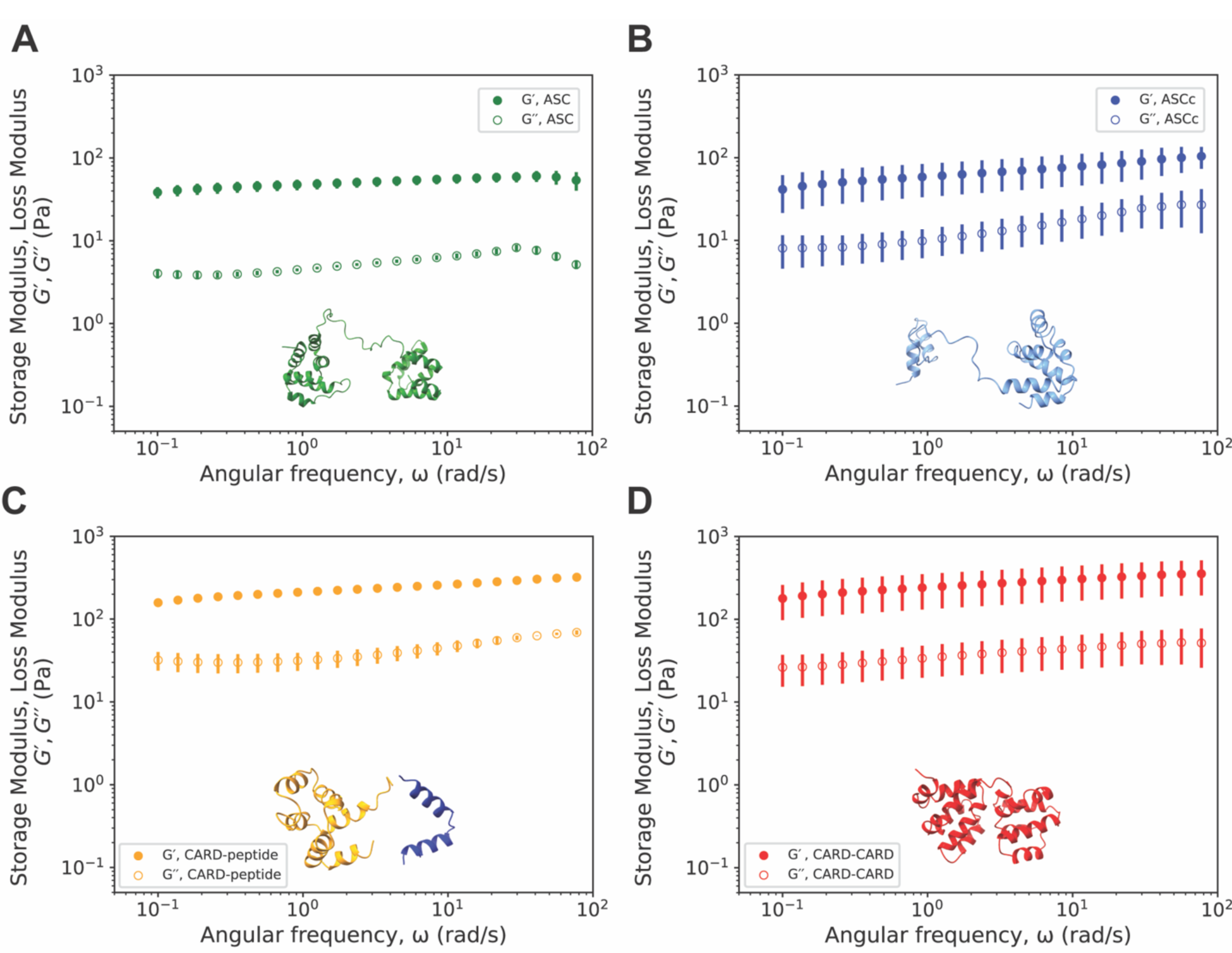
Viscoelastic response of ASC-inspired hydrogels as a function of shear strain. Log-log plot of storage (G’ solid symbols) and loss (G’’ open symbols) moduli as a function of shear strain at constant frequency of 1 Hz (6.28 rad/s) for A) ASC, B) ASCc, C) CARD-peptide, and D) CARD-CARD. Data reported for each condition is the average of two independent measurements. Error bars represent the standard error of the mean.

**Figure 8.**
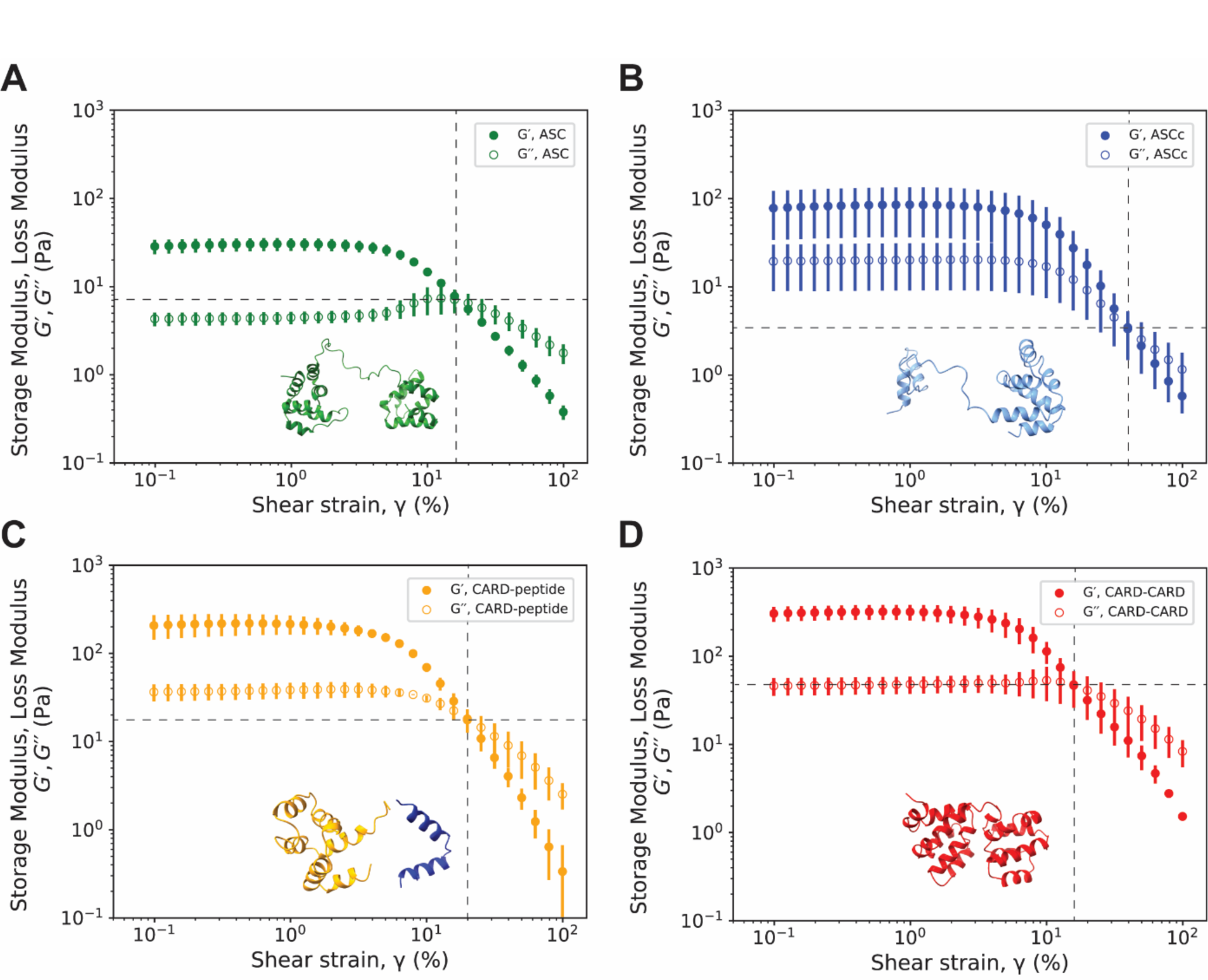
Viscoelastic response of ASC-inspired hydrogels as a function of angular frequency. Log-log plot of storage (G’, solid symbols) and loss (G’’, open symbols) moduli as a function of angular frequency at 1% shear strain for A) ASC, B) ASCc, C) CARD-peptide, and D) CARD-CARD. Data reported for each condition is the average of two independent measurements. Error bars represent the standard error of the mean.

Initial measurements of G’ and G’’ can be assembled in two groups based on similarity (Table 3). One group corresponds to the hydrogels formed by the natural proteins (ASC and ASCc), which have smaller values, and the other group with larger values includes the hydrogels formed by the artificial constructs (CARD-peptide and CARD-CARD) (Table 3). It is worth noting that the difference between G’ and G’’ is significantly smaller for the natural hydrogels than for the designed ones. These results could have biological regulatory consequences, as the natural oligomerization of ASC for inflammasome formation could be activated partly by changes in cytosolic conditions favoring gelation.

**Table 3.**
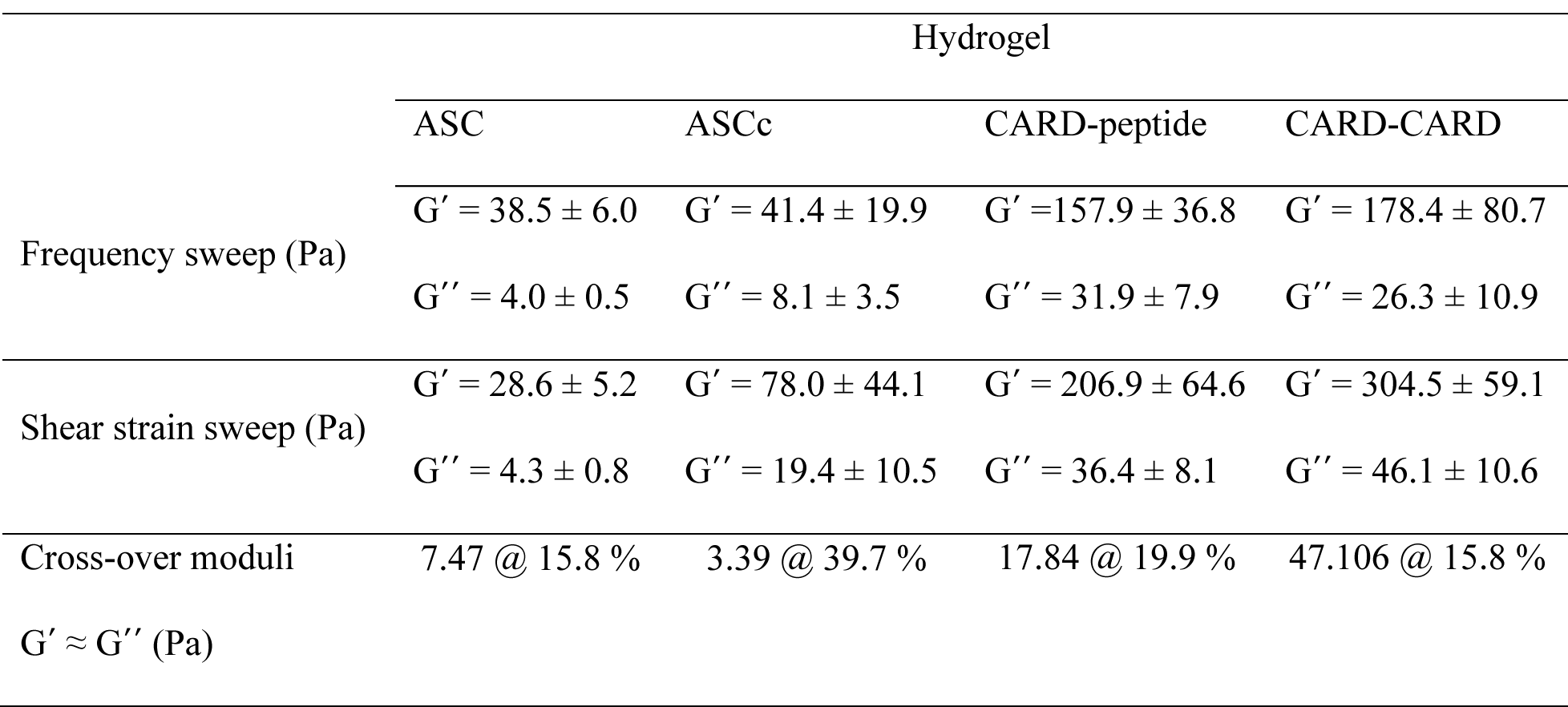
Initial storage (G’) and loss (G’’) moduli, and shear strain at the cross over modulus (G’ ≈.

The hydrogelation processes in ASC-inspired hydrogels depends on cross-linkages formed by physical interactions between the globular domains rather than chemical (covalent) interactions, which are common in non-biological, synthetic hydrogels. The type of cross-linkage has an important impact on the hydrogel mechanical properties; physically cross-linked hydrogels are less resistant to mechanical forces compared to covalently linked hydrogels.^5^ Overall, ASC-inspired hydrogels show viscoelastic behaviors comparable to the positively and negatively charged peptides derived from ASC helices with self-assembling moieties, which show G’ values in the 200 – 400 Pa range.^33^

## CONCLUSION

We have shown here that globular folded proteins and domains of the Death Domain superfamily form non-covalent, pH-responsive hydrogels. Our findings provide one of the very few examples of hydrogels created by the self-assembly of globular proteins and domains in their native conformation without the need to use any proteinaceous or non-proteinaceous cross-linkages. The hydrogelation process of Dead Domains is mainly controlled by the solution pH; however, other factors such as molecular composition and hydrogelation time play important roles. In this work, we have studied the hydrogelation of the Death Domain member ASC, carrying a PYD and a CARD. We have shown that its isoform, ASCc, which bears the CARD but lacks a complete PYD, is also capable of forming hydrogels. We have demonstrated the general hydrogelation capabilities of the Death Domains, PYD and CARD, by forming hydrogels with designed protein constructs based on these domains. Thus, our work opens the door to further studies on the hydrogelation capabilities of other members of the Death Domain superfamily such as DED (Dead Effector Domain) and DD (Dead Domain).

Our characterization of the viscoelastic behavior of ASC-inspired hydrogels reveals, as expected for intrinsic physical cross-linkages, predominantly solid-like mechanical responses. However, the designed CARD-CARD and CARD-peptide hydrogels are stiffer than the natural ASC and ASCc hydrogels. The structural studies by TEM and SEM do not report significant variations between the hydrogels that could explain the different viscoelastic behavior. However, because globular protein domains are the hydrogels building blocks, we speculate whether their specific structure and stability could cause variations in the mechanical properties. Additionally, the linkers connecting the domains of the proteins are of different length (Figure 1), possibly influencing protein stability. For instance, the linker tethering the PYD and CARD in ASC is composed of 23 amino acids, whereas the linker in CARD-CARD is 3 amino acids long. The CARD-peptide does not have a linker as this is a heterotypic hydrogel.

As previously reported, the viscoelastic behavior of globular-protein hydrogels at the molecular level does not only depend on the cross-linkages but on the load-bearing modules (proteins chains) as well.^1,17,23–25^ When globular protein-based hydrogels are subjected to mechanical forces, the cross-linkages are considered the force transducers, whereas the load-bearing modules control the mechanical response.^17^ Mechanical forces can unfold globular proteins, thus, the unfolding/folding processes become critical factors controlling the hydrogel viscoelastic response. The cross-linkages in ASC-inspired hydrogels are formed by entanglement of filaments and fibrils *via* electrostatic and hydrophobic interactions. However, the load-bearing modules are the actual globular domains and the linkers connecting them. Therefore, upon the exertion of mechanical stress, the proteins of ASC-inspired hydrogels with different domains and linkers could follow several unfolding pathways, resulting in variations in energy dissipation and polymer extension, ultimately leading to discrepant viscoelastic behaviors.

Further research is needed to investigate the mechanical unfolding of ASC and ASC-inspired proteins, as well as to determine the extension of the polypeptide chain upon unfolding. These studies will shed light into the different viscoelastic response of ASC-inspired hydrogels. Understanding the molecular bases responsible for the mechanical properties of these hydrogels will be key to design potential biotechnological applications. Importantly, ASC-inspired hydrogels can serve as models to study natural biological hydrogelation processes and to investigate the influence of load-bearing modules (proteins) on the hydrogel mechanical behavior in the absence of artificial cross-linkers.

## ACKNOWLEDGEMENTS

P.D.P., A.W. and E. d. A., acknowledge the National Institute of Allergy and Infectious Diseases of the National Institutes of Health for financial support under award number R15AI146780 to E.d.A. and 3R15AI146780-01S1 to A.W. and E.d.A. P.D.P., A.W., A.M.S., E.G-M, A.G., R.C.A.E and E.d.A. acknowledge support from the NSF-CREST Center for Cellular and Biomolecular Machines at the University of California, Merced (NSF-HRD-1547848 and NSF-HRD-2112675). A.M.S and A.G. acknowledge funding through NSF CBET 2047210. We thank UC Merced Imaging facility for access to the TEM and SEM equipment. We are grateful to Dr. Yue Wang (University of California, Merced) and Dr. Anand Ramasubramanian (San Jose State University) for access to their rheometer instrument. The content of this publication is solely the responsibility of the authors and does not necessarily represent the official views of the National Institutes of Health or the National Science Foundation.

## SUPPORTING INFORMATION

**Supporting Figure 1.**
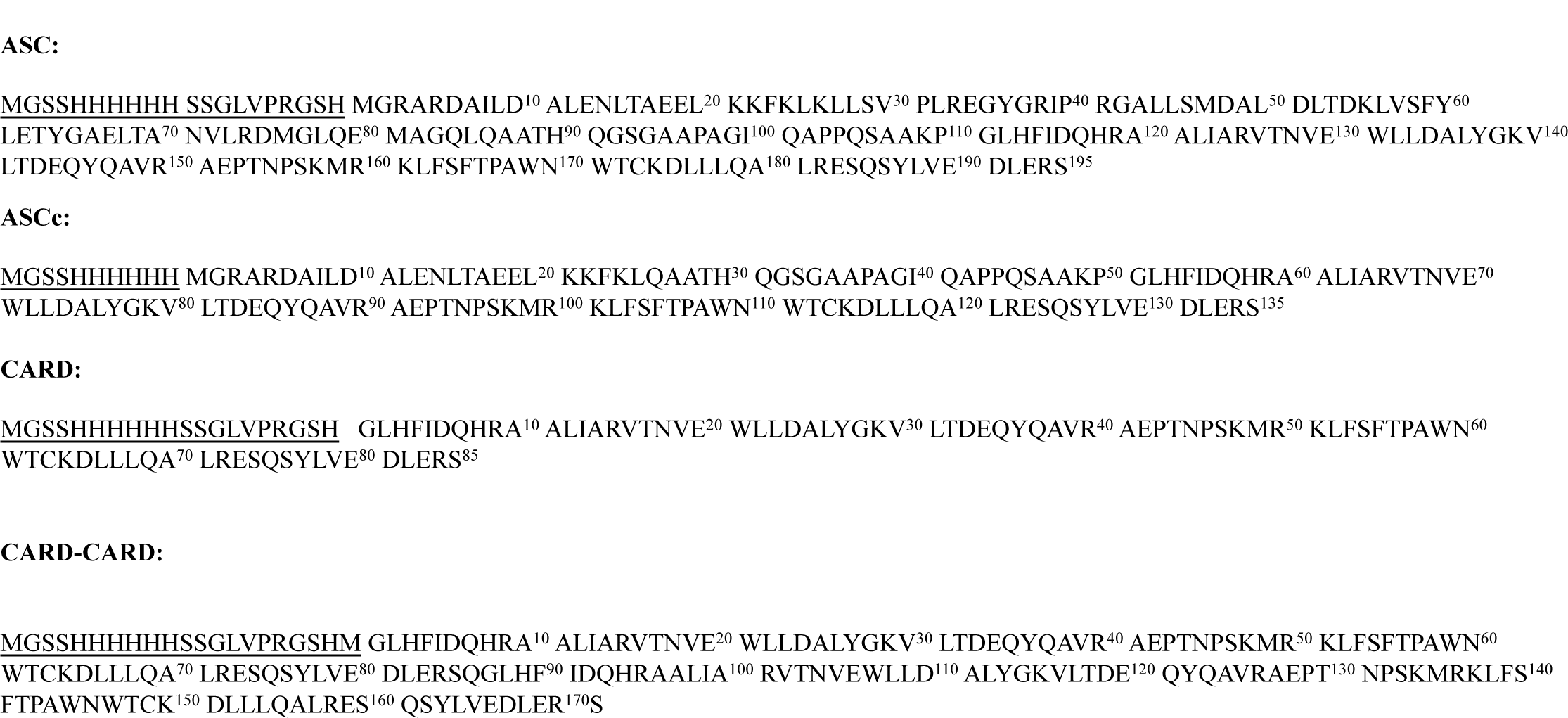
Amino acid sequence of ASC and ASC-derived protein constructs used for hydrogel formation. The amino acid sequences are numbered from N- to C-terminus and the purification tags at the N-terminus are underlined.

**Supporting Figure 2.**
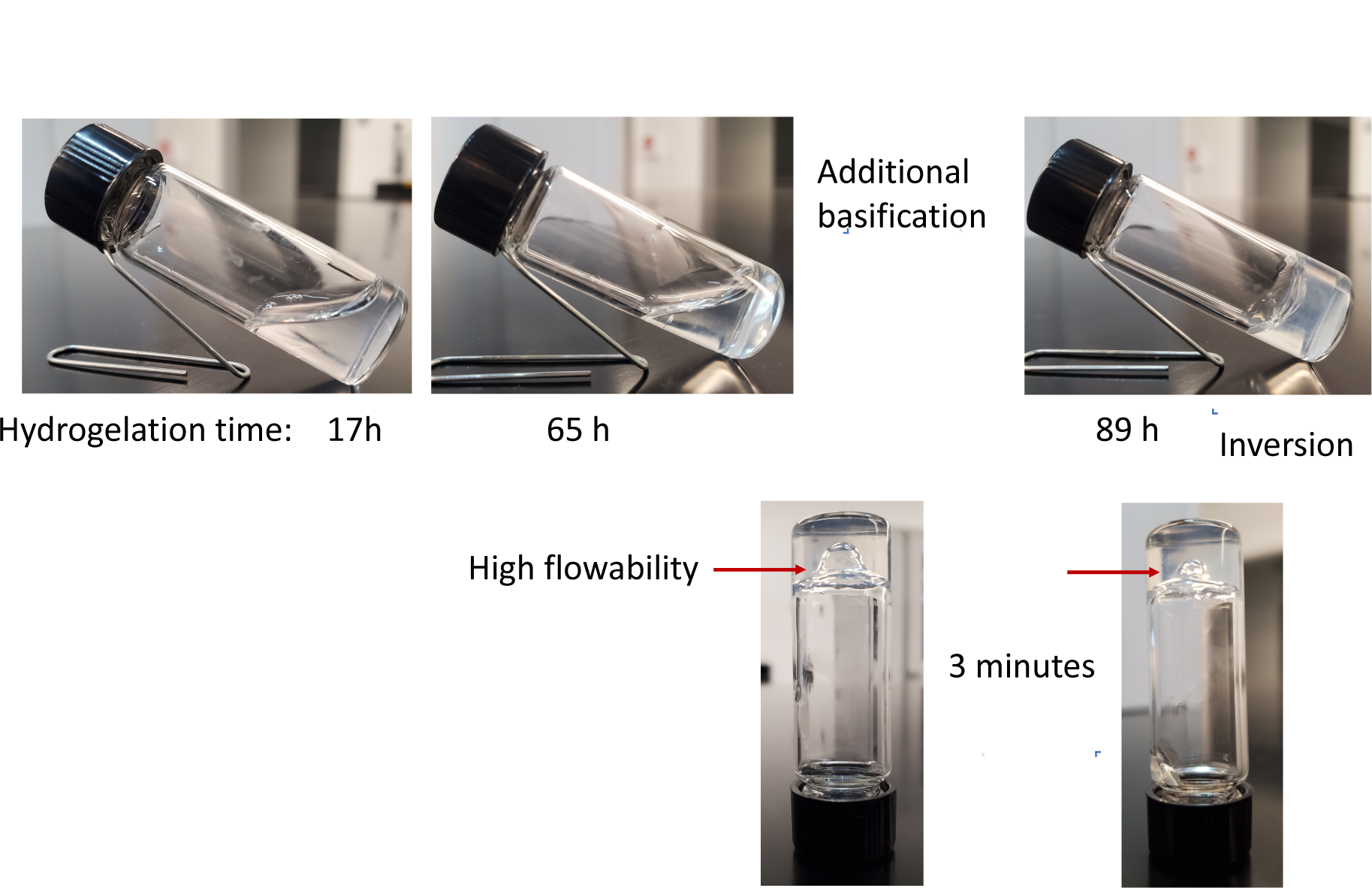
Hydrogelation process of the CARD-only protein. The CARD-only protein needs additional basification and time for hydrogelation. The formed hydrogel shows higher flowability compared to the other hydrogels formed by ASC, ASCc, CARD-peptide and CARD-CARD.

## REFERENCES

(1) Li, Y.; Xue, B.; Cao, Y. 100th Anniversary of Macromolecular Science Viewpoint: Synthetic Protein Hydrogels. ACS Macro Lett 2020, 9 (4), 512–524. https://doi.org/10.1021/acsmacrolett.0c00109.

(2) Yang, T.; Wang, L.; Wu, W.-H.; Wei, S.; Zhang, W.-B. Orchestrating Chemical and Physical Cross-Linking in Protein Hydrogels to Regulate Embryonic Stem Cell Growth. ACS Macro Lett 2023, 12 (2), 269–273. https://doi.org/10.1021/acsmacrolett.2c00741.

(3) Huerta-López, C.; Alegre-Cebollada, J. Protein Hydrogels: The Swiss Army Knife for Enhanced Mechanical and Bioactive Properties of Biomaterials. Nanomaterials 2021, 11 (7). https://doi.org/10.3390/nano11071656.

(4) Ahn, W.; Lee, J.-H.; Kim, S. R.; Lee, J.; Lee, E. J. Designed Protein- and Peptide-Based Hydrogels for Biomedical Sciences. J Mater Chem B 2021, 9 (8), 1919–1940. https://doi.org/10.1039/D0TB02604B.

(5) Jonker, A. M.; Löwik, D. W. P. M.; van Hest, J. C. M. Peptide- and Protein-Based Hydrogels. Chemistry of Materials 2012, 24 (5), 759–773. https://doi.org/10.1021/cm202640w.

(6) Functional Biopolymers; Jafar Mazumder, M. A., Sheardown, H., Al-Ahmed, A., Eds.; Springer International Publishing: Cham, 2019. https://doi.org/10.1007/978-3-319-95990-0.

(7) Mizuguchi, Y.; Mashimo, Y.; Mie, M.; Kobatake, E. Temperature-Responsive Multifunctional Protein Hydrogels with Elastin-like Polypeptides for 3-D Angiogenesis. Biomacromolecules 2020, 21 (3), 1126–1135. https://doi.org/10.1021/acs.biomac.9b01496.

(8) Sun, Y.; Li, W.; Wu, X.; Zhang, N.; Zhang, Y.; Ouyang, S.; Song, X.; Fang, X.; Seeram, R.; Xue, W.; He, L.; Wu, W. Functional Self-Assembling Peptide Nanofiber Hydrogels Designed for Nerve Degeneration. ACS Appl Mater Interfaces 2016, 8 (3), 2348–2359. https://doi.org/10.1021/acsami.5b11473.

(9) Wagner, C. E.; Krupkin, M.; Smith-Dupont, K. B.; Wu, C. M.; Bustos, N. A.; Witten, J.; Ribbeck, K. Comparison of Physicochemical Properties of Native Mucus and Reconstituted Mucin Gels. Biomacromolecules 2023, 24 (2), 628–639. https://doi.org/10.1021/acs.biomac.2c01016.

(10) Miranda-Nieves, D.; Chaikof, E. L. Collagen and Elastin Biomaterials for the Fabrication of Engineered Living Tissues. ACS Biomater Sci Eng 2017, 3 (5), 694–711. https://doi.org/10.1021/acsbiomaterials.6b00250.

(11) Blanco-Fernandez, B.; Ibañez-Fonseca, A.; Orbanic, D.; Ximenes-Carballo, C.; Perez-Amodio, S.; Rodríguez-Cabello, J. C.; Engel, E. Elastin-like Recombinamer Hydrogels as Platforms for Breast Cancer Modeling. Biomacromolecules 2023. https://doi.org/10.1021/acs.biomac.2c01080.

(12) Van Vlierberghe, S.; Dubruel, P.; Schacht, E. Biopolymer-Based Hydrogels as Scaffolds for Tissue Engineering Applications: A Review. Biomacromolecules 2011, 12 (5), 1387–1408. https://doi.org/10.1021/bm200083n.

(13) Banta, S.; Wheeldon, I. R.; Blenner, M. Protein Engineering in the Development of Functional Hydrogels. Annu Rev Biomed Eng 2010, 12 (1), 167–186. https://doi.org/10.1146/annurev-bioeng-070909-105334.

(14) Hume, J.; Sun, J.; Jacquet, R.; Renfrew, P. D.; Martin, J. A.; Bonneau, R.; Gilchrist, M. L.; Montclare, J. K. Engineered Coiled-Coil Protein Microfibers. Biomacromolecules 2014, 15 (10), 3503–3510. https://doi.org/10.1021/bm5004948.

(15) Petka, W. A.; Harden, J. L.; McGrath, K. P.; Wirtz, D.; Tirrell, D. A. Reversible Hydrogels from Self-Assembling Artificial Proteins. Science (1979) 1998, 281 (5375), 389–392. https://doi.org/10.1126/science.281.5375.389.

(16) Haines-Butterick, L.; Rajagopal, K.; Branco, M.; Salick, D.; Rughani, R.; Pilarz, M.; Lamm, M. S.; Pochan, D. J.; Schneider, J. P. Controlling Hydrogelation Kinetics by Peptide Design for Three-Dimensional Encapsulation and Injectable Delivery of Cells. Proceedings of the National Academy of Sciences 2007, 104 (19), 7791–7796. https://doi.org/10.1073/pnas.0701980104.

(17) Wu, J.; Li, P.; Dong, C.; Jiang, H.; Xue, B.; Gao, X.; Qin, M.; Wang, W.; Chen, B.; Cao, Y. Rationally Designed Synthetic Protein Hydrogels with Predictable Mechanical Properties. Nat Commun 2018, 9 (1), 620. https://doi.org/10.1038/s41467-018-02917-6.

(18) Lu, H. D.; Charati, M. B.; Kim, I. L.; Burdick, J. A. Injectable Shear-Thinning Hydrogels Engineered with a Self-Assembling Dock-and-Lock Mechanism. Biomaterials 2012, 33 (7), 2145–2153. https://doi.org/https://doi.org/10.1016/j.biomaterials.2011.11.076.

(19) Fu, L.; Haage, A.; Kong, N.; Tanentzapf, G.; Li, H. Dynamic Protein Hydrogels with Reversibly Tunable Stiffness Regulate Human Lung Fibroblast Spreading Reversibly. Chem. Commun. 2019, 55 (36), 5235–5238. https://doi.org/10.1039/C9CC01276A.

(20) Zakeri, B.; Fierer, J. O.; Celik, E.; Chittock, E. C.; Schwarz-Linek, U.; Moy, V. T.; Howarth, M. Peptide Tag Forming a Rapid Covalent Bond to a Protein, through Engineering a Bacterial Adhesin. Proceedings of the National Academy of Sciences 2012, 109 (12), E690–E697. https://doi.org/10.1073/pnas.1115485109.

(21) Guan, D.; Ramirez, M.; Shao, L.; Jacobsen, D.; Barrera, I.; Lutkenhaus, J.; Chen, Z. Two-Component Protein Hydrogels Assembled Using an Engineered Disulfide-Forming Protein–Ligand Pair. Biomacromolecules 2013, 14 (8), 2909–2916. https://doi.org/10.1021/bm400814u.

(22) Wong Po Foo, C. T. S.; Lee, J. S.; Mulyasasmita, W.; Parisi-Amon, A.; Heilshorn, S. C. Two-Component Protein-Engineered Physical Hydrogels for Cell Encapsulation. Proceedings of the National Academy of Sciences 2009, 106 (52), 22067–22072. https://doi.org/10.1073/pnas.0904851106.

(23) Hanson, B. S.; Dougan, L. Network Growth and Structural Characteristics of Globular Protein Hydrogels. Macromolecules 2020, 53 (17), 7335–7345. https://doi.org/10.1021/acs.macromol.0c00890.

(24) Hughes, M. D. G.; Cussons, S.; Mahmoudi, N.; Brockwell, D. J.; Dougan, L. Single Molecule Protein Stabilisation Translates to Macromolecular Mechanics of a Protein Network. Soft Matter 2020, 16 (27), 6389–6399. https://doi.org/10.1039/C9SM02484K.

(25) Fang, J.; Mehlich, A.; Koga, N.; Huang, J.; Koga, R.; Gao, X.; Hu, C.; Jin, C.; Rief, M.; Kast, J.; Baker, D.; Li, H. Forced Protein Unfolding Leads to Highly Elastic and Tough Protein Hydrogels. Nat Commun 2013, 4 (1), 2974. https://doi.org/10.1038/ncomms3974.

(26) Park, H. H.; Lo, Y.-C.; Lin, S.-C.; Wang, L.; Yang, J. K.; Wu, H. The Death Domain Superfamily in Intracellular Signaling of Apoptosis and Inflammation. Annu Rev Immunol 2007, 25 (1), 561–586. https://doi.org/10.1146/annurev.immunol.25.022106.141656.

(27) de Alba, E. Structure, Interactions and Self-Assembly of ASC-Dependent Inflammasomes. Arch Biochem Biophys 2019, 670, 15–31. https://doi.org/https://doi.org/10.1016/j.abb.2019.05.023.

(28) Diaz-Parga, P.; de Alba, E. Protein Interactions of the Inflammasome Adapter ASC by Solution NMR. Methods Enzymol 2019, 625, 223–252. https://doi.org/10.1016/BS.MIE.2019.07.008.

(29) de Alba, E. Structure and Interdomain Dynamics of Apoptosis-Associated Speck-like Protein Containing a CARD (ASC). Journal of Biological Chemistry 2009, 284 (47), 32932–32941. https://doi.org/10.1074/jbc.M109.024273.

(30) Bryan, N. B.; Dorfleutner, A.; Kramer, S. J.; Yun, C.; Rojanasakul, Y.; Stehlik, C. Differential Splicing of the Apoptosis-Associated Speck like Protein Containing a Caspase Recruitment Domain (ASC) Regulates Inflammasomes. J Inflamm 2010, 7 (1), 23. https://doi.org/10.1186/1476-9255-7-23.

(31) Diaz-Parga, P.; de Alba, E. Inflammasome Regulation by Adaptor Isoforms, ASC and ASCb, via Differential Self-Assembly. Journal of Biological Chemistry 2022, 298 (3), 101566. https://doi.org/https://doi.org/10.1016/j.jbc.2022.101566.

(32) Expasy. https://www.expasy.org.

(33) Shy, A. N.; Wang, H.; Feng, Z.; Xu, B. Heterotypic Supramolecular Hydrogels Formed by Noncovalent Interactions in Inflammasomes. Molecules 2021, 26 (1). https://doi.org/10.3390/molecules26010077.

(34) Nambayan, R. J. T.; Sandin, S. I.; Quint, D. A.; Satyadi, D. M.; de Alba, E. The Inflammasome Adapter ASC Assembles into Filaments with Integral Participation of Its Two Death Domains, PYD and CARD. Journal of Biological Chemistry 2019, 294 (2), 439–452. https://doi.org/https://doi.org/10.1074/jbc.RA118.004407.

(35) Diaz-Parga, P.; Gould, A.; de Alba, E. Natural and Engineered Inflammasome Adapter Proteins Reveal Optimum Linker Length for Self-Assembly. Journal of Biological Chemistry 2022, 298 (11). https://doi.org/10.1016/j.jbc.2022.102501.

(36) Oroz, J.; Barrera-Vilarmau, S.; Alfonso, C.; Rivas, G.; de Alba, E. ASC Pyrin Domain Self-Associates and Binds NLRP3 Protein Using Equivalent Binding Interfaces. Journal of Biological Chemistry 2016, 291 (37), 19487–19501. https://doi.org/https://doi.org/10.1074/jbc.M116.741082.

(37) Sharma, M.; de Alba, E. Structure, Activation and Regulation of NLRP3 and AIM2 Inflammasomes. Int J Mol Sci 2021, 22 (2). https://doi.org/10.3390/ijms22020872.

(38) Wang, H.; Feng, Z.; Lu, A.; Jiang, Y.; Wu, H.; Xu, B. Instant Hydrogelation Inspired by Inflammasomes. Angewandte Chemie International Edition 2017, 56 (26), 7579–7583. https://doi.org/https://doi.org/10.1002/anie.201702783.

(39) Sakakeeny, L.; Roubenoff, R.; Obin, M.; Fontes, J. D.; Benjamin, E. J.; Bujanover, Y.; Jacques, P. F.; Selhub, J. Plasma Pyridoxal-5-Phosphate Is Inversely Associated with Systemic Markers of Inflammation in a Population of U. S. Adults. J Nutr 2012, 142 (7), 1280–1285. https://doi.org/https://doi.org/10.3945/jn.111.153056.

(40) Yang, J.; Zhang, Y. I-TASSER Server: New Development for Protein Structure and Function Predictions. Nucleic Acids Res 2015, 43 (W1), W174–W181. https://doi.org/10.1093/nar/gkv342.

(41) Pettersen, E. F.; Goddard, T. D.; Huang, C. C.; Meng, E. C.; Couch, G. S.; Croll, T. I.; Morris, J. H.; Ferrin, T. E. UCSF ChimeraX: Structure Visualization for Researchers, Educators, and Developers. Protein Science 2021, 30 (1), 70–82. https://doi.org/https://doi.org/10.1002/pro.3943.

